# Proliferation Saturation Index in an adaptive Bayesian approach to predict patient-specific radiotherapy responses

**DOI:** 10.1101/469957

**Authors:** Enakshi D. Sunassee, Dean Tan, Tianlin Ji, Renee Brady, Eduardo G. Moros, Jimmy J. Caudell, Slav Yartsev, Heiko Enderling

**Author notes:** Correspondence: Heiko Enderling, Ph.D., Department of Integrated Mathematical Oncology, H. Lee Moffitt Cancer Center & Research, Institute, Tampa, FL 33612, USA.

## Abstract

**Purpose:** Radiotherapy prescription dose and dose fractionation protocols vary little between individual patients having the same tumor grade and stage. To personalize radiotherapy a predictive model is needed to simulate radiation response. Previous modeling attempts with multiple variables and parameters have been shown to yield excellent data fits at the cost of nonidentifiability and clinically unrealistic results.

**Materials and Methods:** We develop a mathematical model based on a proliferation saturation index (PSI) that is a measurement of pre-treatment tumor volume-to-carrying capacity ratio that modulates intrinsic tumor growth and radiation response rates. In an adaptive Bayesian approach, we utilize an increasing number of data points for individual patients for predicting response to subsequent radiation doses.

**Results:** Model analysis shows that using PSI as the only patient-specific parameter, model simulations can fit longitudinal clinical data with high accuracy (R^2^=0.84). By analyzing tumor response to radiation using daily CT scans early in the treatment, response to the remaining treatment fractions can be predicted after two weeks with high accuracy (c-index=0.89).

**Conclusion:** The PSI model may be suited to forecast treatment response for individual patients and offer actionable decision points for mid-treatment protocol adaptation. The presented work provides an actionable image-derived biomarker prior to and during therapy to personalize and adapt radiotherapy.

## Introduction

Radiation therapy (RT) is the single most utilized therapeutic agent in oncology (Torres-Roca 2012). Radiation is generally delivered in many small fractions to achieve a large total dose. Fractionation of radiotherapy provides temporal windows for normal tissue recovery, and conventional radiation is delivered in fraction sizes of 1.8 to 2.0 Gy once a day, Monday through Friday. Efforts to adapt RT mid-treatment due to anatomy changes and radiographic response mainly address adapting the target volume to limit toxicity to adjacent organs-at-risk (Hansen et al. 2006; Woodford et al. 2007). Predicting which patient will respond with tumor regression significant for treatment adaptation and when this will occur is of high clinical value.

Mathematical and physical modeling has a long history in radiation therapy, including the linear-quadratic dose-dependent survival probability concept (LQ-Model) (Fowler 1989), biologically equivalent dose calculations (Fowler 2010), tumor cure probability estimations (O’Rourke et al. 2009) and normal tissue complication probability analysis (Brodin et al. 2018). Inroads have been made more recently into using patient-specific data to simulate untreated tumor growth, predict how individual patients would response to radiation, and how to evaluate clinical responses (Rockne et al. 2010; Corwin et al. 2013; Prokopiou et al. 2015; Poleszczuk et al. 2017).

In a recent modeling study, a variety of tumor growth and treatment response models with different biological resolutions and complexities were analyzed against clinical data of non-small cell lung cancer (Tariq et al. 2016). An adaptive Bayesian approach was introduced to derive patient-specific model parameters with increasing availability of longitudinal volumetric response data early in radiation therapy to predict volumetric regression during the remainder of the treatment. This study used the data of twenty five non-small cell lung cancer (NSCLC) patients previously reported (Woodford et al. 2007). The tumor volume was available for each patient at all radiation fractionation time points. With such large data set, the used Akaike Information Criterion (AIC) analysis selected for models with better fits to the data despite larger degrees of freedom (number of parameters) (Burnham & Anderson 2002). The chosen model was a two-variable, coupled differential equation model simulating total tumor volume as the sum of living and dead cancer cells (*V=V_l_+V_d_*). To best fit the model to the data, parameters were derived that would kill all living cancer cells after 4-6 Gy in 2-3 days (of the conventional 45 days schedule), and the modeled exponential decay of the dead cell compartment mimicked the overall tumor volume regression (Tariq et al. 2016). Whilst mathematically elegant with excellent fits to the clinical data, the model implications are clinically unrealistic and yield to predictions that cannot be trusted. We have previously developed a single-variable, radiotherapy response model that simulates patient-specific volume regression during radiation using the concept of a proliferation saturation index, PSI (Prokopiou et al. 2015; Poleszczuk et al. 2017). We propose the PSI model to simulate the discussed clinical data with less biological resolution (one-variable, one parameter equation) but high predictive power.

Tumors are composites of proliferating and hypoxic growth arrested cells. Overall radiation response depends on their respective proportions at irradiation. In multi-compartment mathematical models that distinguish between proliferating and growth-arrested cells, proliferation and oxygenation status-dependent radiation response can be simulated on the cellular level (Enderling et al. 2009; Gao et al. 2013). Tumor growth *in vivo* can be approximated by Gompertzian or logistic growth dynamics (Gerlee 2013; Enderling & Chaplain 2014; Benzekry et al. 2014). Initial exponential growth at low cell densities when most cells have access to ample resources decelerates when cells at the core of the tumor become growth-arrested, mainly due to limited space and exhausted intratumoral nutrient supply as resources are consumed by cells closer to the tumor surface. This established the notion of a tumor carrying capacity (K) as the maximum tumor volume that can be supported by a given environment. A tumor carrying capacity may evolve depending on the oxygen and nutrients supply through tissue vascularization (Hahnfeldt et al. 1999), removal of metabolic waste products (Freyer & Sutherland 1986), and evasion of immune surveillance (Dunn et al. 2002). Hence, the ratio of the initial pre-treatment tumor volume to its carrying capacity ratio (V_0_/K) describes the saturation in overall tumor cell proliferation as the tumor approaches its carrying capacity, which we call PSI (Prokopiou et al. 2015). At treatment start, PSI=V_0_/K, reflects the history of the co-evolution of the tumor with its environment – and thus it can be assumed to provide patient/tumor-specific information. Tumor volumes close to their carrying capacity, i.e. a high PSI, have only a small proportion of proliferating cells that are most sensitive to radiation-induced damage.

## Materials and Methods

### Mathematical model

We propose logistic growth of tumor volume increasing proportional to the intrinsic potential doubling time, λ = 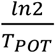day^−1^, scaled by the current environmental condition-dependent carrying capacity (*K*):

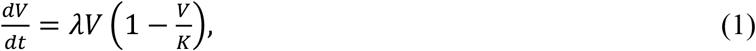

where *V* is the tumor volume at time *t*, and *dV/dt* is the change of tumor volume over time. Radiation response is modeled as instantaneous volume changes *V_PostIR_* = *V_preIR_* − 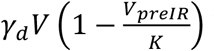 before and after each radiation fraction, where *γ_d_* = 1 − e^−(*αd+βd^2^*)^ represents radiation-induced death after dose *d* following the LQ-model with radiosensitivity parameters *α*(Gy^−1^) and *β* (Gy^−2^). It follows from eqn. (1) that larger *V/K* ratios present in patients with a high PSI lead to small volume changes and low fraction of proliferating cell in radiation-resistant tumors, whereas tumors with low PSI have more proliferative cells and thus are more radiosensitive. In this model, two patients that present with similar tumor volumes could have a different PSI, which would result in different responses to the same radiotherapy protocol. To predict patient-specific radiation response, tumor carrying capacity *K* and thus PSI before radiation needs to be estimated.

### Patient population

Twenty-five non-small cell lung cancer patients were treated for NSCLC with standard of care radiation treatment over an 8-week period on the Hi-Art Helical Tomotherapy unit at the London Regional Cancer Program, Ontario, Canada, from 2005 to 2007 (Tariq et al. 2016). Volumetric tumor measurements were collected at each treatment fraction.

### Data fitting and parameter estimation

In line with Tariq’s work and references within (Tariq et al. 2016; Jin et al. 2010), we set *α/β*=10 Gy for NSCLC. We utilize a genetic algorithm to derive parameter values for which the sum of residuals between data and simulation results is minimized (Prokopiou et al. 2015). We first find the best parameter values for each individual patient. Then we find the best tumor growth and radiation sensitivity parameters that are uniform across all patients with only PSI being patient-specific. To compare the 3-parameter and 1-parameter models, we calculate the AIC using

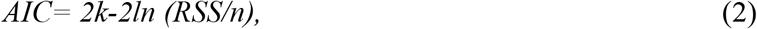

where *k* is the number of model parameters, *n* is the number of data samples, and *RRS* is the residual sum of squares between model simulation and data.

### Adaptive Bayesian Prediction

The twenty-five patients were randomized into model training (13 patients) and prediction (12 patients) cohorts. Growth and radiation response parameters are derived for patients in the model training cohort, that are then kept constant for patients in the prediction cohort. For treatment response prediction, data points from the first treatment week are utilized to estimate patient-specific PSI using derived growth rate and radiation response parameters and tumor carrying capacity, *K*. From these, radiation response is simulated for the remainder of treatment. PSI is recalculated and treatment response prediction is re-evaluated after each subsequent treatment week using all patient data up to that time point. A concordance C-index is calculated as a measure of the difference between the model-predicted and clinically observed tumor volumes for each patient to evaluate the prediction performance of the mathematical model (Harrell et al. 1982; Walker et al. 2017). A C-index=0.5 represents poor predictive ability (random guessing), and as the c-index approaches 1.0 it represents a very strong predictive ability of the model.

## Results

### PSI model fits the data with individual patient-specific parameters

The PSI model with parameter sets *[λ_i_, α_i_, PSI_i_]* fits the data for each individual patient *i* with high accuracy (overall R^2^=0.91, 0.0-0.98; **Fig. 1A**). Median tumor growth rate was *λ=0.01* day^−1^ (mean 0.07; 0.002-0.99), median radiation sensitivity was *α=0.06* (mean 0.08; 0.013-0.312), and median PSI was PSI=0.88 (mean 0.7; 0.05-0.999; **Fig. 1B**). As in the Tariq study (Tariq et al. 2016) the best fit to individual patient’s data was achieved for patient 9 (R^2^=0.98, **Fig. 1C**). A patient with average mean squared error between model and data (R^2^=0.81) is shown in **Fig. 1D**.

**Figure 1.**
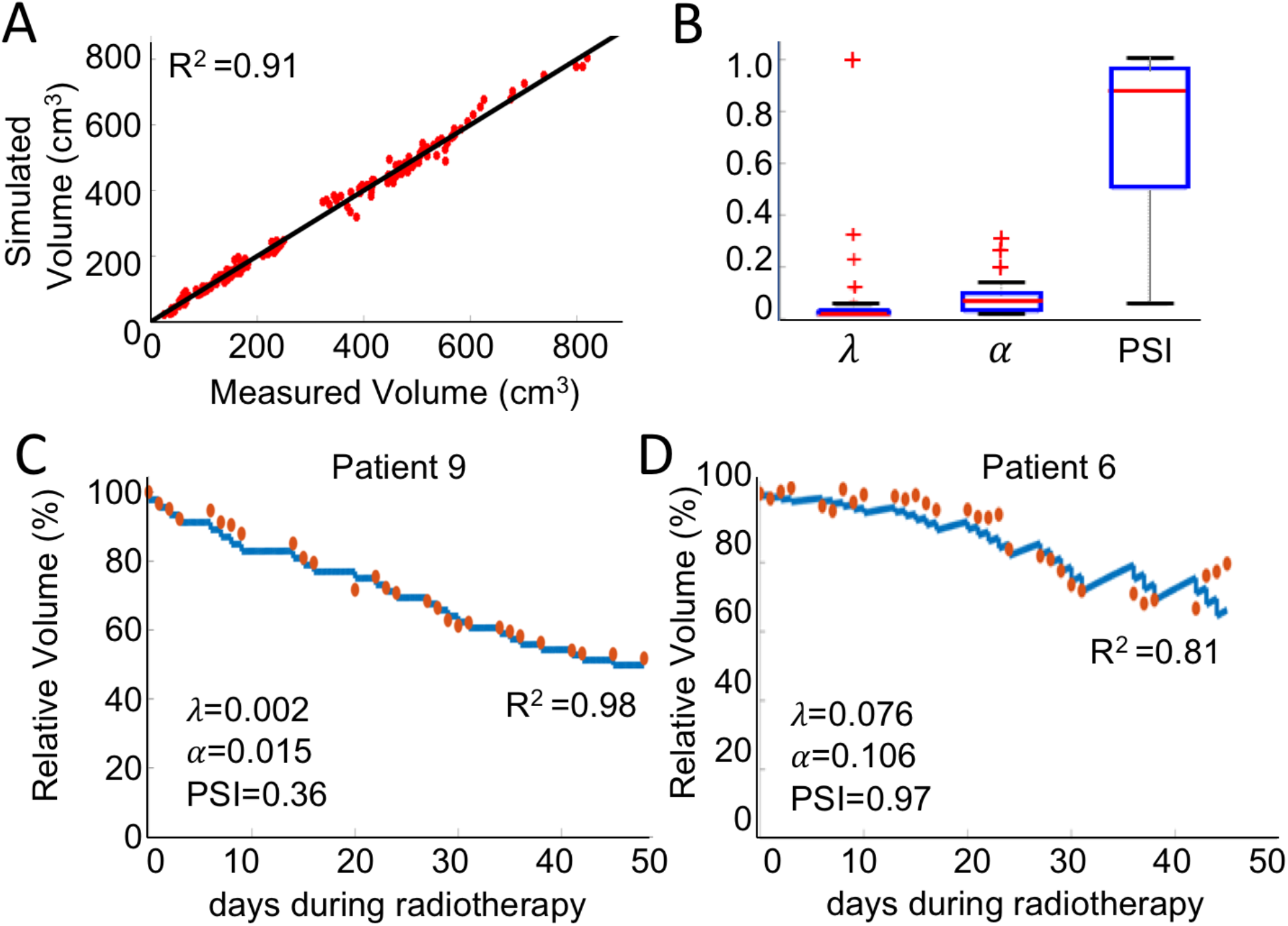
Simulated patient-specific tumor volume regression during radiotherapy. **A.** Correlation of simulated volumes and measured volumes for all 25 patients (R^2^=0.91). **B.** Distribution of model parameter values across all 25 patients. **C.** Patient 9 with the best fit of the mathematical model (blue curve) to the data (red circles, R^2^=0.98). **D.** Patient 6 with an average fit of the model to the data (R^2^=0.81).

### PSI model fits the data with PSI as single patient-specific parameter

The standard deviation of the distribution of tumor growth rate *λ* and radiation sensitivity a are significantly smaller than the deviation of the PSI distribution (coefficient of variation CoV=150%; **Fig. 1B**). This suggests that PSI may be contributing more to patient-specific treatment responses, and that tumor growth rate *λ* and radiation sensitivity *α* may be kept uniform across all patients. Simultaneous fit of the model to all patient data yields uniform tumor growth parameter *λ=0.012* day^−1^ and radiation sensitivity *α=0.029* Gy^−1^ with only patient-specific PSI. This one-parameter model slightly reduces quality of fit (R^2^=0.84; **Fig. 2A**) but yields a significantly lower median AIC (−4.5 vs. 0.45, p=0.00017). As model parameters are non-orthogonal, keeping l and a constant reduces the coefficient of variation for the distribution of PSI across all patients compared to the three-parameter model (CoV=46% vs. CoV=150%; **Fig. 2B**). Patient 9 with the best three-parameter model fit remains the best fit of the 1-parameter model to the data with the goodness of fit only dropping one percentage point (R^2^=0.97; **Fig. 2C**). Patient 6 remains an average-fit patient 6 (R^2^=0.77; **Fig. 2D**).

**Figure 2.**
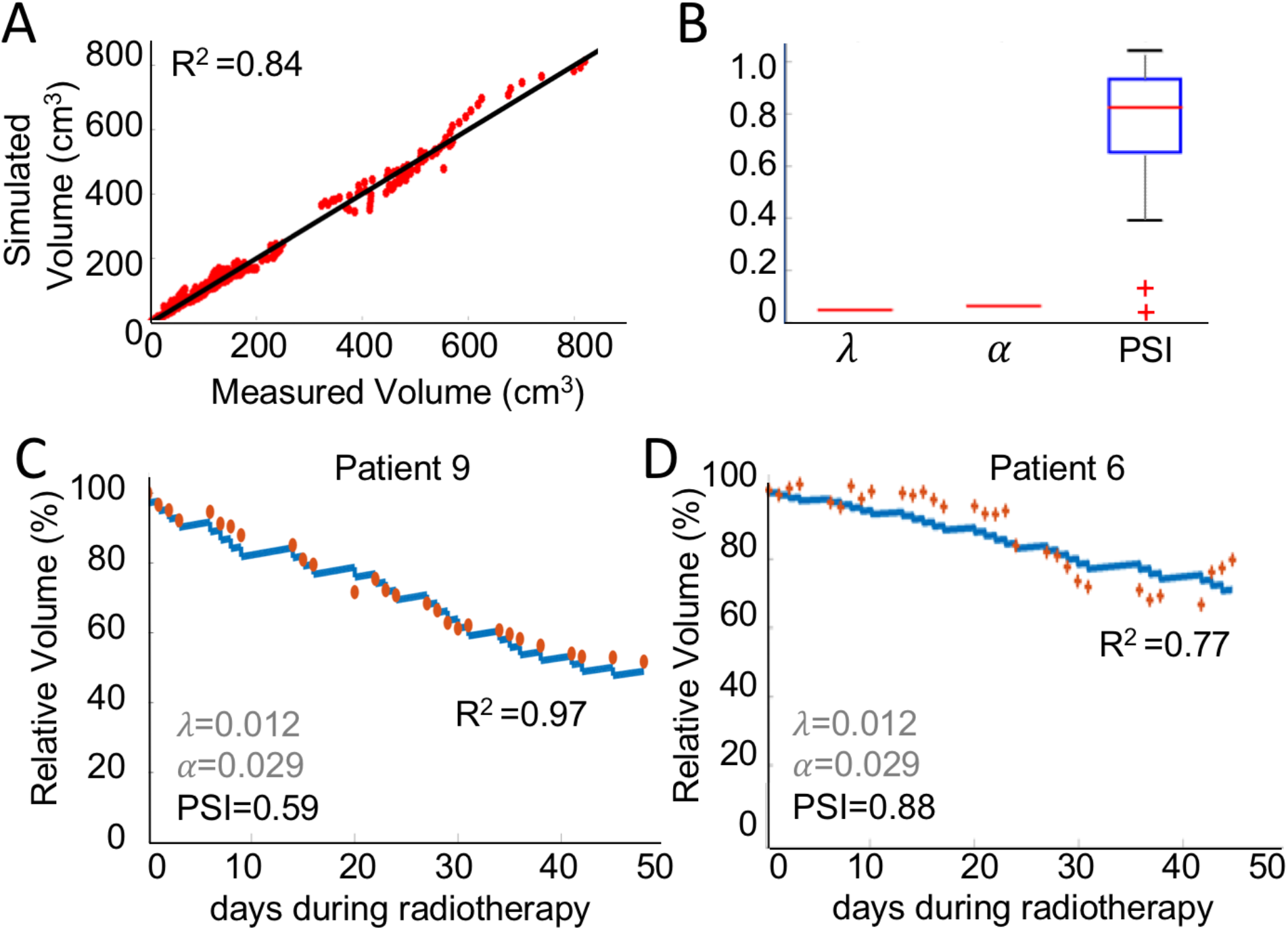
Simulated patient-specific tumor volume regression during radiotherapy with only PSI as patient-specific parameter with derived uniform tumor growth parameter *λ=0.012* day^−1^ and radiation sensitivity *α=0.029* Gy^−1^. **A.** Correlation of simulated volumes and measured volumes for all 25 patients (R^2^=0.84). **B.** Distribution of PSI across all 25 patients. **C.** Patient 9 with the best fit of the mathematical model to the data (R^2^=0.97). **D.** Patient 6 with an average fit of the model to the data (R^2^=0.77).

### Adaptive Bayesian prediction of treatment responses

Figure 3 shows Bayesian prediction plots for sample patients with varying quality of prediction performances. (C-index). Figure 3D shows the variation of C-indices dependent on increasing patient-specific data used for PSI calculation for predictions. With patient-specific tumor volumes of one treatment week and growth and death parameters from the model training cohort, the median C-index>0.86 (0.67-0.98) is significantly restricted indicating an accuracy in prediction of NSCLC response to radiotherapy as soon as week 2 of therapy.

**Figure 3.**
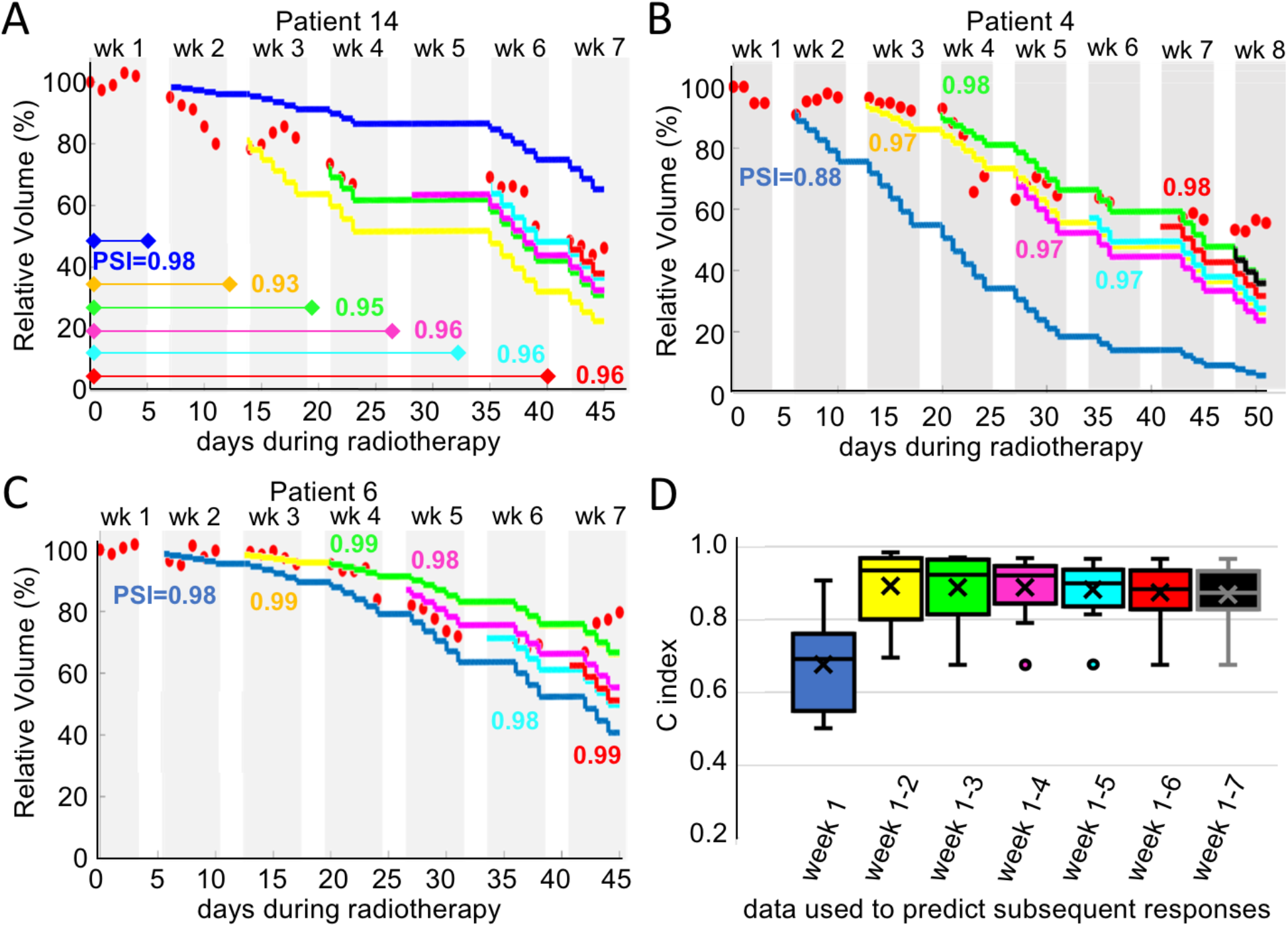
Adaptive Bayesian prediction of radiation response. Model training derived parameters are *λ=0.001* day^−1^ and *α=0.069* Gy^−1^. Red circles: patient data. Colored curves: Model-predicted trajectories. Colored numbers: Derived PSI for the data up to marked time point. **A.** Patient 14 with high prediction accuracy after 2 weeks data. **B.** Patient 4 with median prediction accuracy. **C**. Patient 6 with lowest prediction accuracy. Using just four weeks of data, subsequent responses can be predicted with high accuracy. **D.** Distribution of C-index for prediction accuracy for all 25 patients as a function of number of weeks used to derive patient-specific PSIs.

## Discussion

Mathematical modeling is becoming an emerging tool in simulating clinical oncology data. Whilst the field is still in its infancy, most work is focused on analyzing retrospective data to build mathematical models and evaluate their predictive potential for patient-specific treatment responses (Rockne et al. 2010; Yankeelov et al. 2015; Geng et al. 2017). The art of the trade is to balance and synergize available data, model complexity and model usefulness. From limited resolution of clinical data reconstruction of radiobiological parameters may be an ill-posed problem (Chvetsov et al. 2015). With introduction of additional variables that have not and cannot be identified longitudinally in a noninvasive manner should be met with caution as it may render the model non-identifiable and generate clinically unrealistic predictions (Chvetsov et al. 2014; Tariq et al. 2016).

Here we have proposed to use a simple mathematical model of tumor growth and radiation therapy response that has one patient-specific parameter. The proliferation saturation index, PSI, a measure of the volume-to-carrying capacity ratio in individual patients, is undoubtedly a gross oversimplification of the complex biology underlying tumor growth and response to radiation. However, despite its simplicity or maybe because of its simplicity, the PSI model may be able to derive patient-specific cancer and cancer response properties that could be predictive and prognostic. We set out to re-evaluate the radiation response of twenty-five NSCLC patients with the simple PSI model (Prokopiou et al. 2015) and compare its adaptive Bayesian prediction power to the more complex model introduced by Tariq (Tariq et al. 2016). In our simple logistic growth and radiation response model, the model parameters tumor growth rate, carrying capacity, and radiation sensitivity are not independent. Changing one of the three parameters could yield identical fits to longitudinal patient data after adaptation of the other two parameters. By analyzing model fits, we identified the tumor carrying capacity, and in particular the tumor volume-to-carrying capacity ratio, as the most likely driver of interpatient heterogeneity in radiotherapy responses. The error introduced by a uniform tumor growth rate and radiosensitivity for all patients is kept small as both parameters are multiplied by a factor containing the remaining lose parameter, the patient-specific carrying capacity. Despite a slight reduction in goodness of fit, fixing the growth and death parameters enables higher confidence in calculation of PSI and ultimately higher predictive power. The presented work is retrospective in nature and limited to a previously published dataset in NSCLC. Before general conclusions can be drawn about the applicability of the PSI framework for NSCLC or other tumors treated with radiation, validation in an independent cohort is of utmost importance.

The model assumed that the underlying dynamics follow logistic growth. Different growth laws may provide equally good fits of to the data but may predict grossly different responses to therapy (Poleszczuk et al. 2017). Therefore, as mathematical oncologists we are well advised to develop, compare, and compete multiple models with different assumptions and complexities against each other to evaluate where model predictions disagree or agree with one another. This may raise confidence in in silico predictions and have highest chances of success for clinical translation.

## Acknowledgements

This project was supported in part by a pilot award from the NIH/NCI U54CA143970-05 (Physical Science Oncology Network (PSON)) “Cancer as a complex adaptive system”. DT and TJ were supported by the High school Internship Program in Integrated Mathematical Oncology (HIP IMO) at Moffitt Cancer Center.

